# Bacterial Normalized Binary Fission Growth Model

**DOI:** 10.1101/2021.10.05.463248

**Authors:** Alan Baer, John P. Keady

**Affiliations:** National Center for Biodefense and Infectious Diseases, George Mason University, 10650 Pyramid Place, Manassas, VA, 20110; Department of Physics and Astronomy, GMU, 4400 University Drive, MSN 6A2, Fairfax, VA, 22030, Hearing Ergonomics and Acoustic Resources LLC, Fairfax Station, VA, 22039

**Keywords:** Bacterial growth model, Optical density curve, Acoustic disruption, Bacterial inhibition, Bacterial growth curves, Escherichia coli, bacterial size

## Abstract

Mathematical models have traditionally been used to facilitate the interpretation of bacterial growth curves in order to more accurately understand and identify variations in bacterial proliferation. Here, a binary fission growth model was developed to normalize starting bacterial levels, allowing for the identification of changes in bacterial growth and the separation of a bacterial population as it correlates to size. This normalized binary fission model (NBF) relies on a multi-bin growth mode, where each bin is associated with a size range during a growth cycle. The proposed NBF model allows for a determination of the percentage of treated bacteria eradicated compared to a control sample, either generally across all bacterial binary fission sizes or specific to a size range or bin. Comparisons between the NBF model and experimental observations demonstrates that bacterial growth curves, and the ratio of sample growth to a control, can be used to both determine and normalize initial variations in bacterial size, and quantity, among test samples, as well as identify final nutrient levels and the percentage of bacteria affected by treatment.

**Significance Statement:** It is difficult to determine the effectiveness of selective bacterial eradication based upon a bacteria’s characteristics size, related to acoustic resonance. Here we develop a binary fission model to analyze effect on growth curves of size depend eradications. Understanding the effect on growth curves provides a method to extract the eradication percentage and initial bacterial level differences between treated sample and control sample, by using the ratio of treated growth curve to control growth curve.

## Introduction

Bacterial binary fission occurs when a bacterium (e.g., E.coli) elongates during growth until separation occurs, creating two distinct bacteria. Interestingly during binary fission, while a bacterium undergoes elongation across its lengths, a near constant cross sectional dimension is maintained. Many prior models have sought to simulate various aspects of bacterial growth and these analytical models have increasingly been applied to bacterial growth data in an effort to more accurately interpret growth characteristics across sample sets (Rickett et al., 2015). Rickett et. Al. 2015 seeks to use Bayesian statistics to detect differences between bacterial growth rates, using a 4 parameter Baranyi and Roberts model (Baranyi et al., 1993, Baranyi et al., 1994, Rickett et al., 2015). Huang seeks to model, the lag, exponential and transition phases, using exponential and logarithmic functions and compares the developed model to the Baranyi and Roberts Model (Huang, 2008, Huang 2010). However, in each of the previously mentioned models, growth is considered uniform and the bacteria is treated as a homogenous population. In the novel normalized binary fission (NBF) model presented here, growing bacteria is treated as a heterogenous population, with distinct subsets comprising binned populations which are separated according to size range. In the NBF model, each bin includes a population of bacteria (here represented by E. coli) by size range, comprising separately binned populations. As the population in each bin gradually increases, it then spills over and populates the next bin, until reaching the final maximum sized bin. After the final bin size undergoes binary fission, the lowest bin is subsequently increased in population and modeling is continued as desired over the course of bacterial growth. The model is used to examine the effect of various percentages of bacterial eradication on ratio curves to identify regions of growth curves that can be used to extract eradication percentage.

Analyzing bacterial growth typically relies on either direct methods, counting (traditionally cumbersome), or through indirect methods, such as measuring turbidity through the optical density (OD) of bacterial suspensions in comparison to a blank/media only control. High throughput bacterial sampling often relies on indirect OD testing using multi-well plate formats (e.g., 96-Well Plate), were large sample sets of bacteria, along with standards and controls and can easily be jointly monitored. However, when using indirect measurement methods for low volume samples (as occurs during high throughput monitoring), minor variations in bacterial sample starting numbers and media volumes can result in significant analytical challenges when comparing sample growth curves. Even the most precise sample dispensation techniques can result in minor variations in initial bacterial levels and/or nutrient broth levels. Thus, the binary fission model developed here examines simulated growth curves and growth curve ratios for various initial bacterial levels and nutrient broth levels, and develops a corrective methodology, allowing for variations in initial bacterial and nutrient broth across sample sets. Additionally, the binary growth model developed is used to examine the effect of various eradication percentages of treated bacteria on the growth curve, resulting in the development of a method of determining the eradication percentage as a result of treatment compared to a control sample.

Growth curves typically undergo a lag, exponential, and stationary phase. Figure 1 illustrates a classical growth model developed by Baranyi and Roberts (Baranyi et al., 1993, Baranyi et al., 1994, Ricket et al., 2015) illustrating E. coli growth from binary fission. With most bacteria, including E.coli, size is most often directly correlated with the timing of fission. As E.coli size changes during binary fission, it typically grows in one dimension (longitudinal), while remaining essentially unchanged in the axial (width) dimension. Hence at any given time a heterogenous range of bacterium sizes occur, from a minimum size just after fission to a maximum size just prior to fission.

**Figure. 1.**
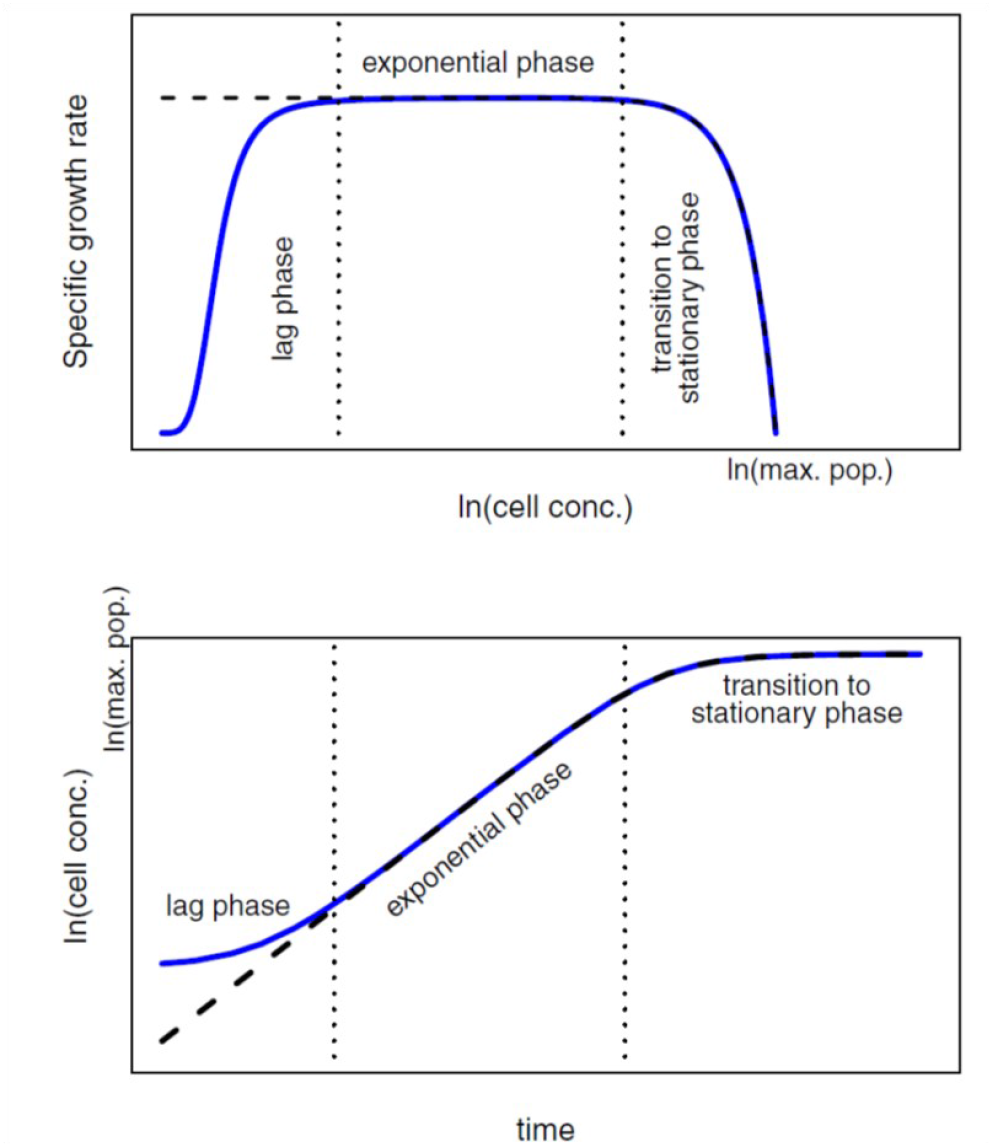
An illustration of the three stages of bacterial growth (lag, exponential and stationary) described in the model of Baranyi and Roberts [Rickett et al., 2015] (solid blue line). Classical models for the population size, *x(t)*, incorporate the exponential and stationary phases (black dashed line) on the logarithmic scale. The top panel shows how the specific growth rate changes during the three stages, whilst the bottom panel demonstrates how these phases correspond to the growth curves observed over time

During growth, the width of E-coli remains impressively stable and is about 1.26 μm ± 0.16 μm (Volkmer et al., 2011). For example, stationary (i.e. in a starved no growth condition) E. coli strain BW25113, has an average length of 1.6 μm ± 0.4 μm, a width of 1.26 μm ± 0.16 μm, and a volume of 1.5μm^3^ ± 1.2 μm^3^. While the same strain of E. coli, in a growth medium of Luria Broth (exponential phase) has an average length of 3.9 μm ± 0.9 μm, a width of 1.26 μm ± 0.16 μm, and a volume of 4.4 μm^3^ ± 1.1 μm^3^ (Volkmer et al., 2011).

The difficulty with current growth models is their inability to extract bacterial eradication data from turbidity data (Optical Density, OD) and to account for differences in initial bacterial levels between the samples and controls, as well as the difference in final nutrient levels between samples and controls. In addition, most current growth models do not directly extract eradication percentage from growth curves, nor do they provide a realizable method for identifying initial differences in bacterial levels between samples and controls. The NBF model proposed here, provides a method of determining the percentage of bacteria eradicated during treatment, informs if the treatment is size specific, and provides a means for identifying and correcting curves for initial bacterial differences (nutrient and bacterial numbers). The NBF model described herein, examines bacterial eradication effects on growth curves and presents a method of using a sample/control growth curve to extract the percentage of bacteria eradicated by various treatments, in addition to a method for adjusting unequal initial bacterial concentrations between controls and samples and nutrient differences.

## Discussion: Method

### Normalized Binary Fission (NBF) Model

In the Normalized (sample/control) Binary Fission Model (NBF Model), it is assumed that the bacteria undergo growth changes in elongation length where the largest elongation length occurs just prior to binary fission. Here, the full size range of bacteria, from the smallest dimension to the largest, are broken into bins, each having a population, and each associated with a size range. For example, Figure 2 illustrates a 9 bin model using a length dimension of 3.9μm ± 0.9μm (Volkmer et al., 2011) with a bandwidth 0.2 μm, where the red dashes represent bacteria.

**Figure. 2.**
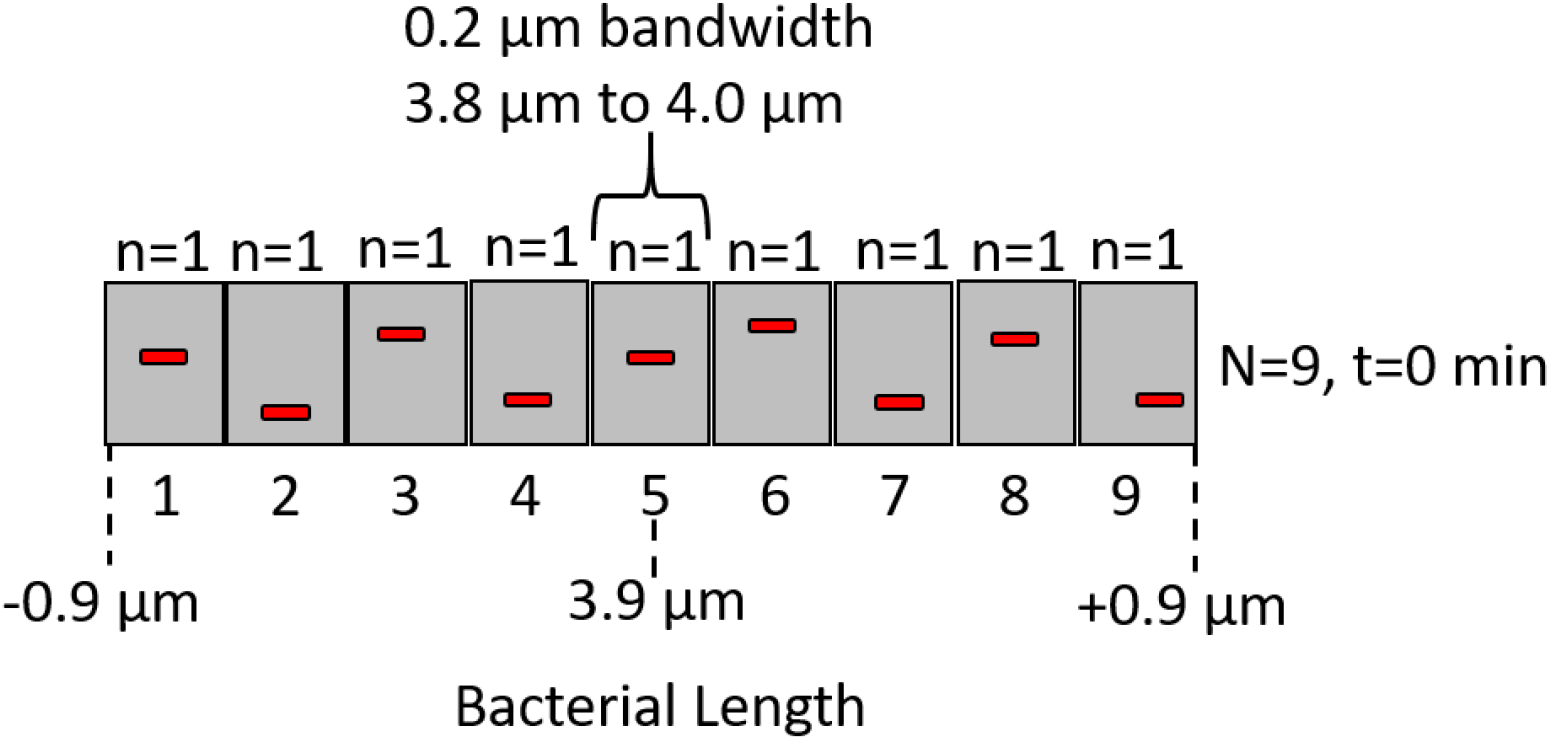
A bin-model with 9 bins, each associated with a particular length bandwidth of bacteria, each bin populated by a single bacteria (red)

In this model, is assumed that the bin associated with the largest size, immediately prior to binary fission, is depopulated following its movement beyond its maximum size, with the lowest sized bin subsequently accepting its population as it divides. Figure 3 illustrates a 25 bin model over two minutes, indicating population of the lowest length related bin. Note that at T=1 minute, the lowest bin contains 2N0 due to each of the largest size bacteria at binary fission breaking into two small bacteria.

**Figure. 3.**
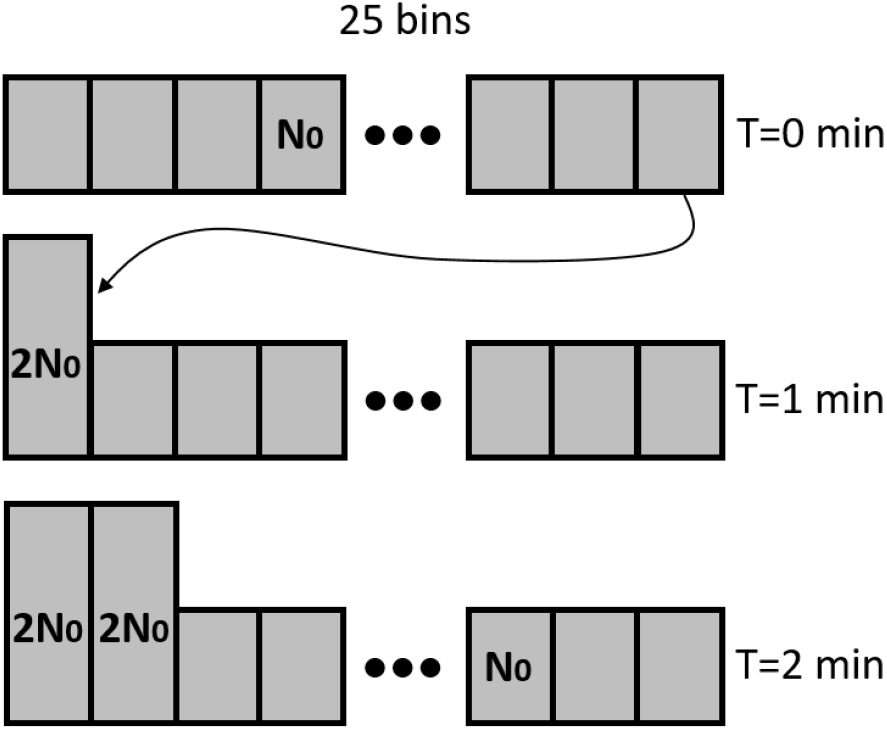
A bin-model with 25 bins, each associated with a particular length bandwidth of bacteria, each bin initially populated by N0 bacteria in each bin, and the growth of the bin population over two minutes, the dots representing the intermediate 18 bins.

Binary fission population growth can be expressed as:

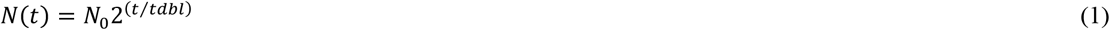

Where N(t) is the number of bacteria at time t, N_0_ is the number of bacteria initially, and the time to double the concentration (tdbl) is the time step upon which binary fission occurs. For example, if the time step is 25 minutes to replicate (tdbl=25min) and the initial population in the lowest bin is N_0_=4, after two time steps (t=2tdbl=50minutes) the population will be N(2tdbl)=16. Note that eqn. (1) is the formula for a particular bin in which the population in that particular bin increases. When examining a treated sample N_t_(t) with a control sample N_c_(t), often the initial bacterial levels N_ot_ and N_oc_ can determine the growth level at a particular time. If both the control and treated samples are grown identically the treated bacterial number at a given time can be expressed as:

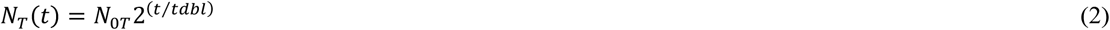

Where N_T_(t) is the remaining (i.e. after treatment) bacterial number at time t and N_0T_ is the initial bacterial amount to be treated (i.e. at the start of treatment) at time 0sec. The control sample can be expressed as:

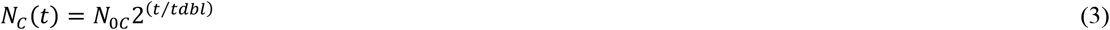

Where N_C_(t) is the control bacterial number at time t and N_0C_ is the initial bacterial amount of the control at time 0sec.

Examining the ratio of treated to control population, eqn. (2)/eqn. (3) one obtains the ratio of initial bacterial levels of the treated bacterial sample to the control sample, expressed as:

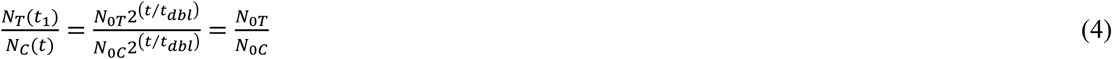

Hence, the ratio of the bacterial population of treated sample to control at t=0, represents the ratio of different live bacterial start levels. This can be used to correct growth curves by adjusting the time dependent treated sample curve by dividing each value by the ratio of eqn. (4).

The final bacterial growth levels depend on available nutrients. The remaining available food (nutrient level) can be expressed as, in terms of the number of bacteria that can be produced, F(t) in a sample at time t is a function of the initial nutrient level (F_0_) and the number of bacteria N(t) which have consumed nutrients so far, which can be expressed as:

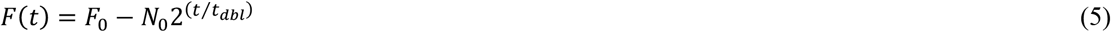

Eqn. (5) expresses the food (nutrient) level in terms of increment units consumed by the bacteria, where a unit nutrient needed is the amount consumed by a bacterium from its smallest size, until it undergoes binary fission.

For example, F_0_=3 refers to a nutrient level sufficient for three bacterium to form. Solving for the time, t_l_, when the available nutrient expires (F(t_l_)=0) one obtains:

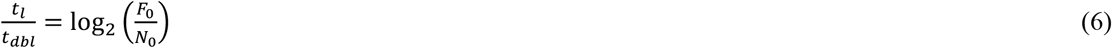

Solving for the bacterial number N at t_l_ we obtain:

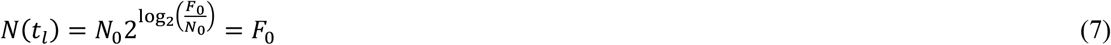

Note that the final bacterial number N(t_l_) is independent of the start bacterial number N_0_ and is dependent only upon the available nutrient level, F_0_. Hence bacterial growth curves, assuming no death, plateau in the stationary phase at the bacterial level that grows in complete consumption of F_0_. The ratio of treated to control population at t=t_l_, using eqn. (7), is related to the ratio of available nutrient levels (F_0S_ and F_0C_) and can be expressed as:

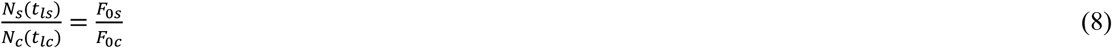

Without correction by the ratio of eqn. (4), the ratio of the bacterial population of treated sample to a control during the stationary phase represents the ratio of the available treated to control nutrient levels. After correction, during the exponential phase, the ratio of the bacterial population of treated sample to control plateaus at the ratio amount of live bacteria between the treated sample and control, providing a method of determining the eradicated level of bacteria from the growth curves. Figure 4 illustrates a growth curve and a ratio of growth curves indicating the relative regions of the curves relevant to eqn. (4) and eqn. (8).

**Figure. 4.**
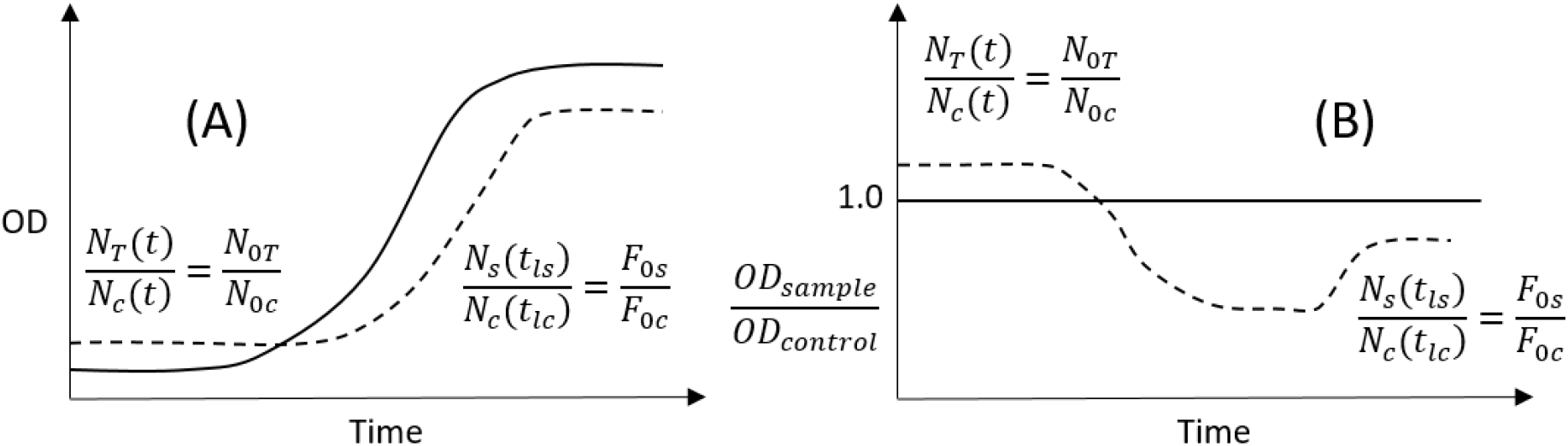
Simulated Optical Density (bacterial number, N) using 600nm light for E.coli, of a treated sample (solid line) and control sample (dashed line) as a function of time (A) and the ratio of Optical Densities (B) of control to control (solid line) and treated to control (dashed line) and the relevant regions of the curves related to equations (4) and (8).

### Model Simulations

Control vs experimental bacterial growths can now be examined. There are two treatment conditions examined, the treatment condition where the bacteria in each bin is affected equally, referred to as linear eradication, and localized eradication where specific bins (sizes) are affected. An example of a linear eradication may be an antibacterial that affects the membrane of E. coli irrespective of bacterial size, and an example of bin specific eradication may be that of a toxin or treatment that disrupts a specific portion of the bacterial life cycle. Figure 5 illustrates simulated Bacterial growth curves for a control (red solid line) and several treated samples in a 25 bin model, using linear eradication. Several eradication levels are illustrated, 12% (red dashed line), 20% (purple dashed line), 40% (light blue dashed line), 60% (dark blue dashed line) and 80% (yellow dashed line) of bacteria eradicated. The simulation generating Figure 5 assumes equal initial bacterial levels for the control and eradicated samples and assumes that the final nutrient levels are also identical. Note that even though the eradicated samples have lagged exponential phases, the final bacterial numbers are identical in time since the nutrient levels are assumed to be identical. Figure 6 illustrates the equivalent growth curve ratios of treated samples to control of Figure 5. Note that the curve ratio plateaus in the exponential phase at the relative remaining live bacterial levels. For example, at 80% the initial bacterial level eradication results in the curve-ratio (green-yellow dashed line) plateauing at a curve-ratio value of 20% indicating that compared to the control the 80% eradication has at 20%. The smoothness of the curve in the lag and exponential phases is a result of linear eradication across bins.

**Figure. 5.**
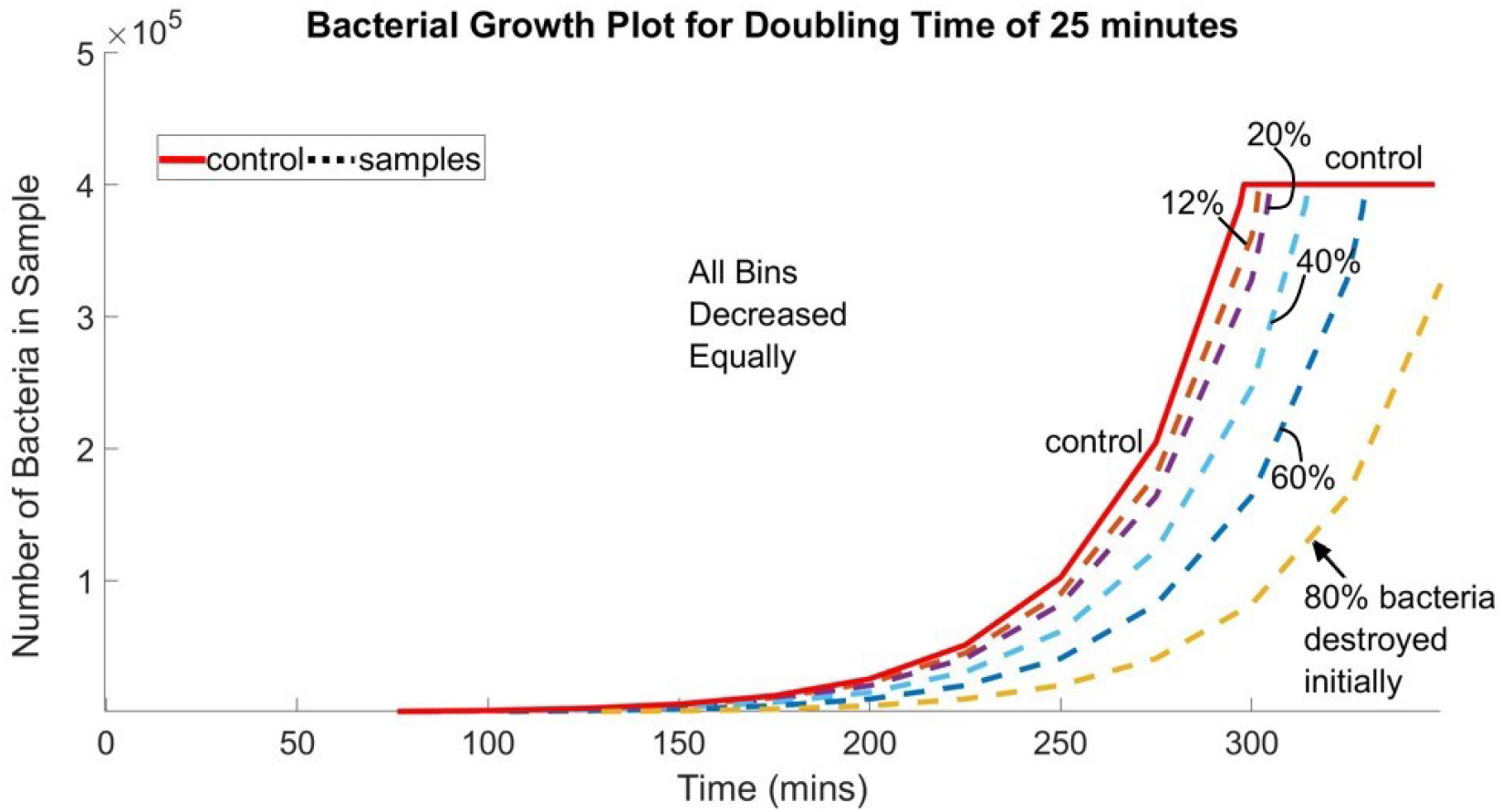
Simulated growth curve values for various linear eradication levels of initial bacteria, with the initial bacterial levels equal and the final food levels the same.

**Figure. 6.**
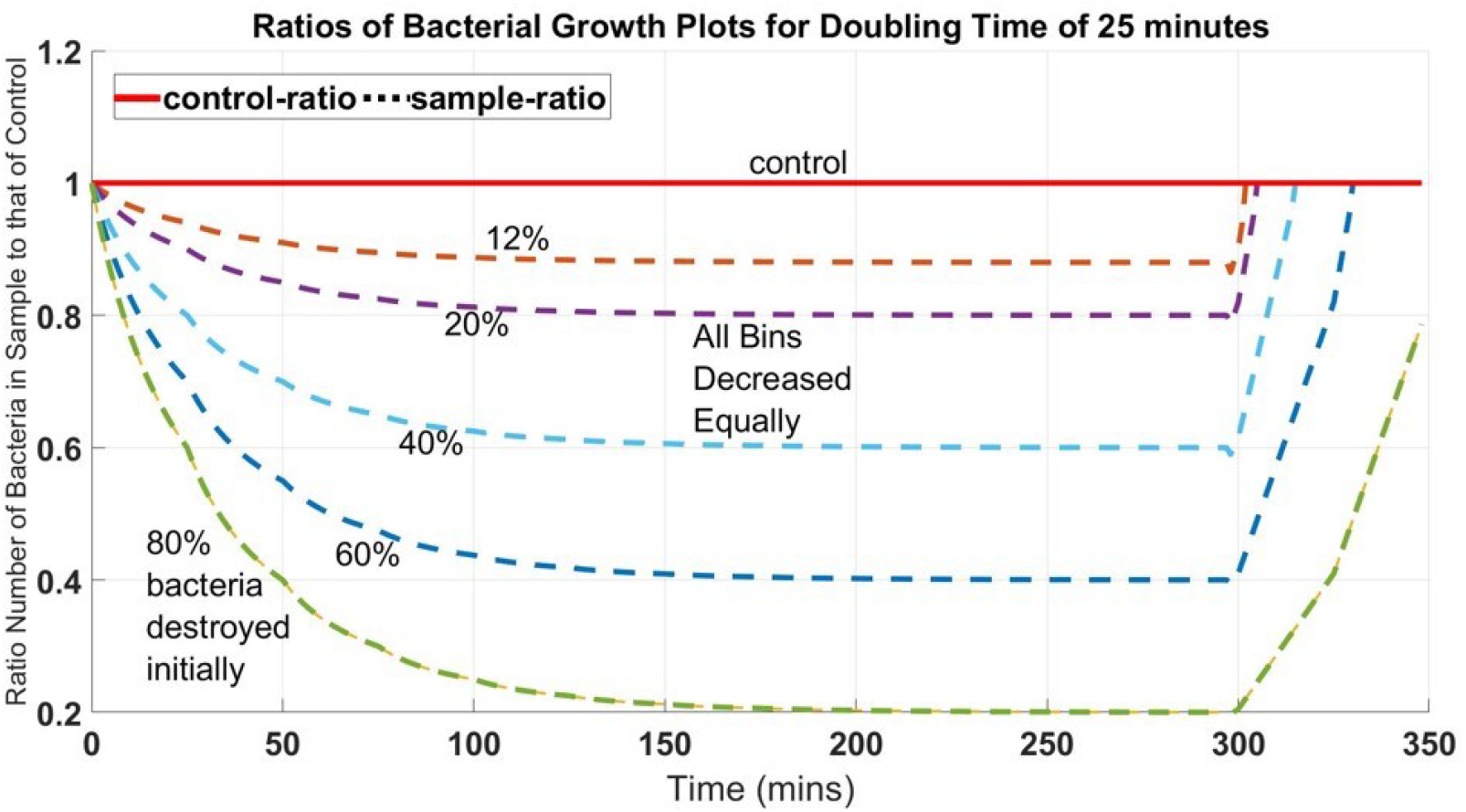
Simulated growth curve-ratio values for the various eradication levels of Figure 5

Figures 5 and 6 simulate eradication across all bin (sizes) equally, Figures 7 and 8 represent the simulated results when eradication of bacteria is size specific, for example only mid bin eradication. Note that Figures 7 and 8 represent conditions where certain bins have been 100% eradicated, although compared to the entire population, which includes the bins not targeted, the total eradication percentage is less than 100%. It is the less than 100% that is reported herein (i.e. the total eradication). Figure 7 is the growth curve of middle bin eradication, where there is 100% eradication in these bins, while Figure 8 is the curve-ratio plots for the same mid bin eradication. Figures 7 and 8 simulate the 20% and 40% eradication curves of Figures 5 and 6 respectively, where the simulated curves of Figures 7 and 8 are based upon a model where the central bins are eradicated first then expanded to either side bins to fulfill the eradication percentage. The stepped nature of the growth curves of Figure 7 are the result of bacteria in mid sizes being eradicated, so that as bacterial grows toward binary fission, the bacterial in the eradicated bins are vacant. Hence, during the period of time when the vacant bin would replicate there is no net bacterial number growth, resulting in a step for the period of time it takes from the next lower populated bin to replicate.

**Figure. 7.**
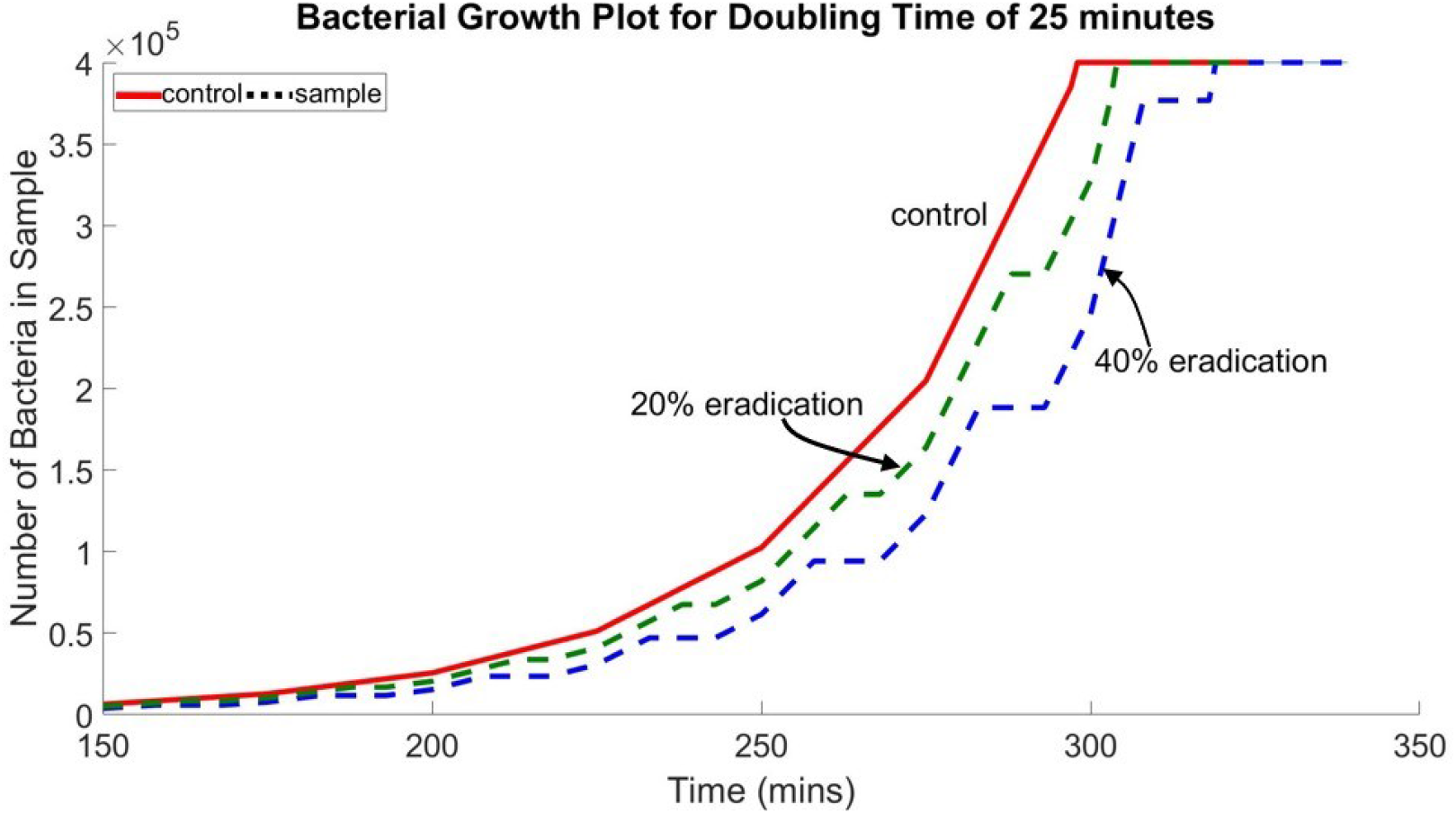
Simulated growth curve values for various mid bin eradication levels of initial bacteria, with the initial bacterial levels equal and the final food levels the same.

**Figure. 8.**
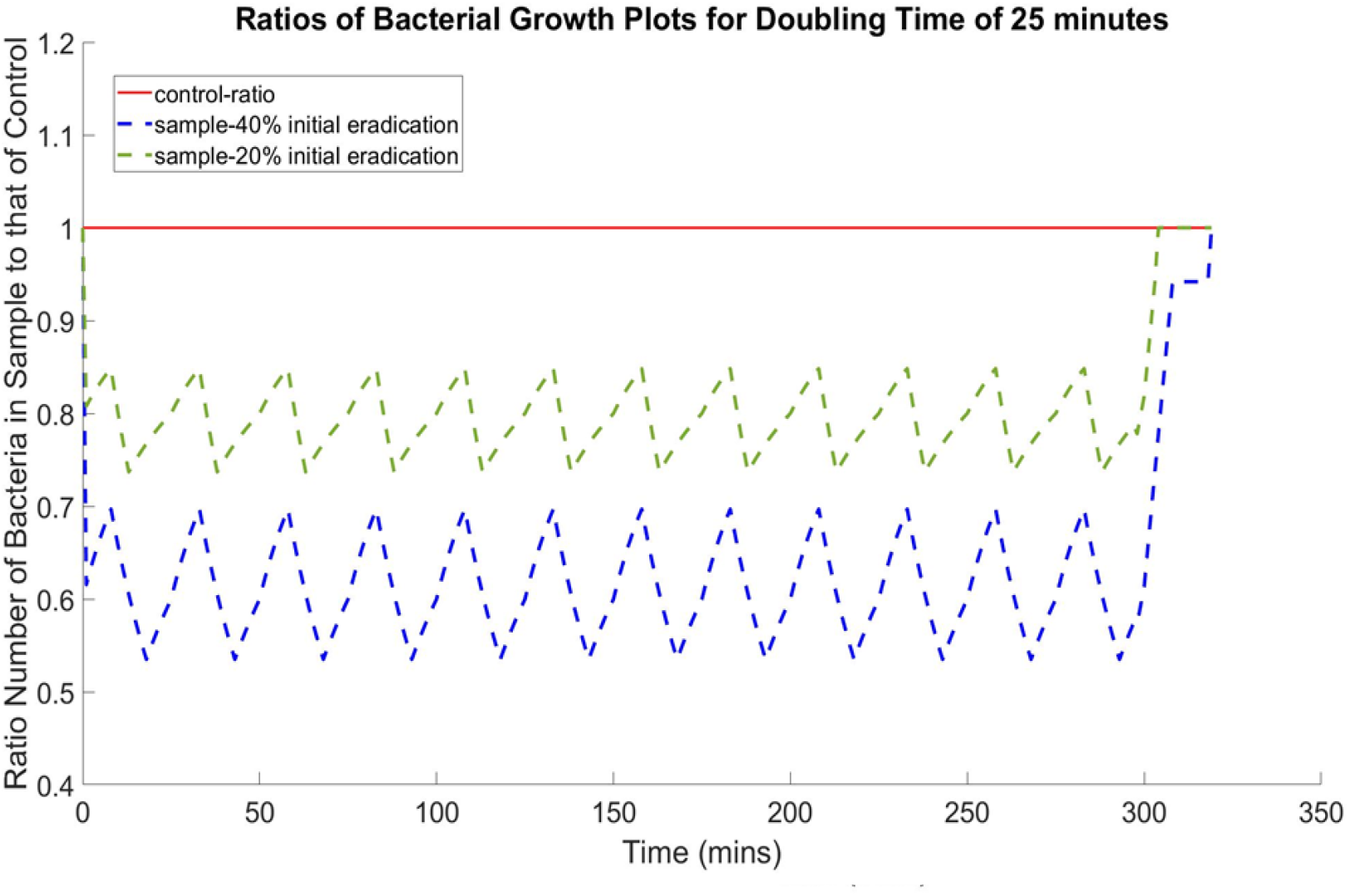
Simulated growth curve-ratio values for the various eradication levels of Figure 7

The step in the growth curve of Figure 7 is replicated as a saw tooth formation in the curve-ratio plots of Figure 8, and if the initial bacterial levels of the sample and control are identical then the saw tooth oscillates at about the eradication level of the sample compared to the control. If the initial bacterial levels of the control are different than the sample prior to treatment, for example slight variations in sample dispensation, the ratio curves of Figure 8 would not start at the same value at time of 0 seconds.

Figures 9 and 10 illustrate the simulated situation where the initial bacterial level of the sample is 110% of the control bacterial numbers, and the final nutrients levels is 90% of the control level.

**Figure. 9.**
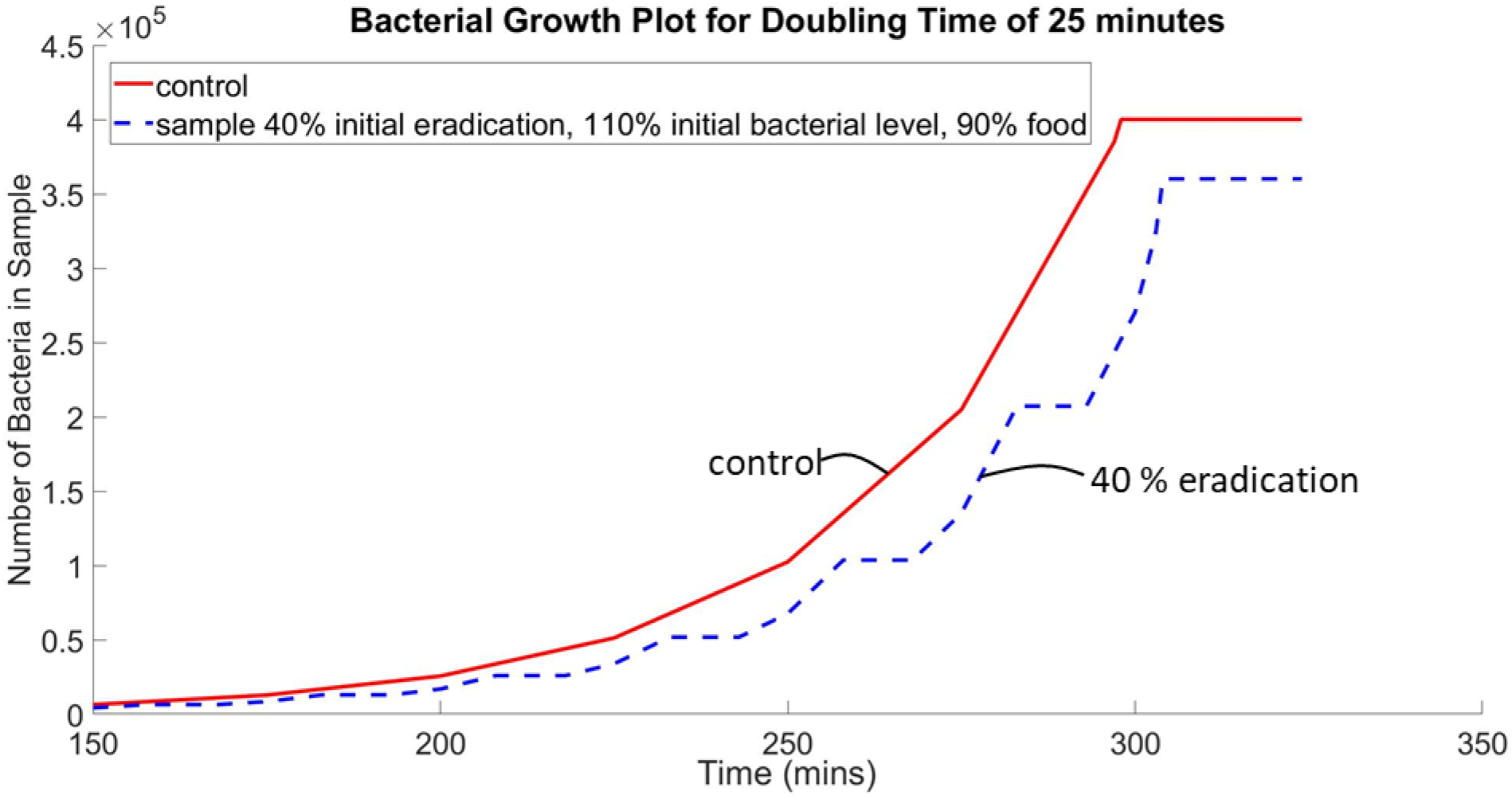
Simulated growth curve values for various mid bin eradication levels of initial bacteria, with the initial bacterial levels of the sample is 110% and the final food levels are 90% of the control values.

**Figure. 10.**
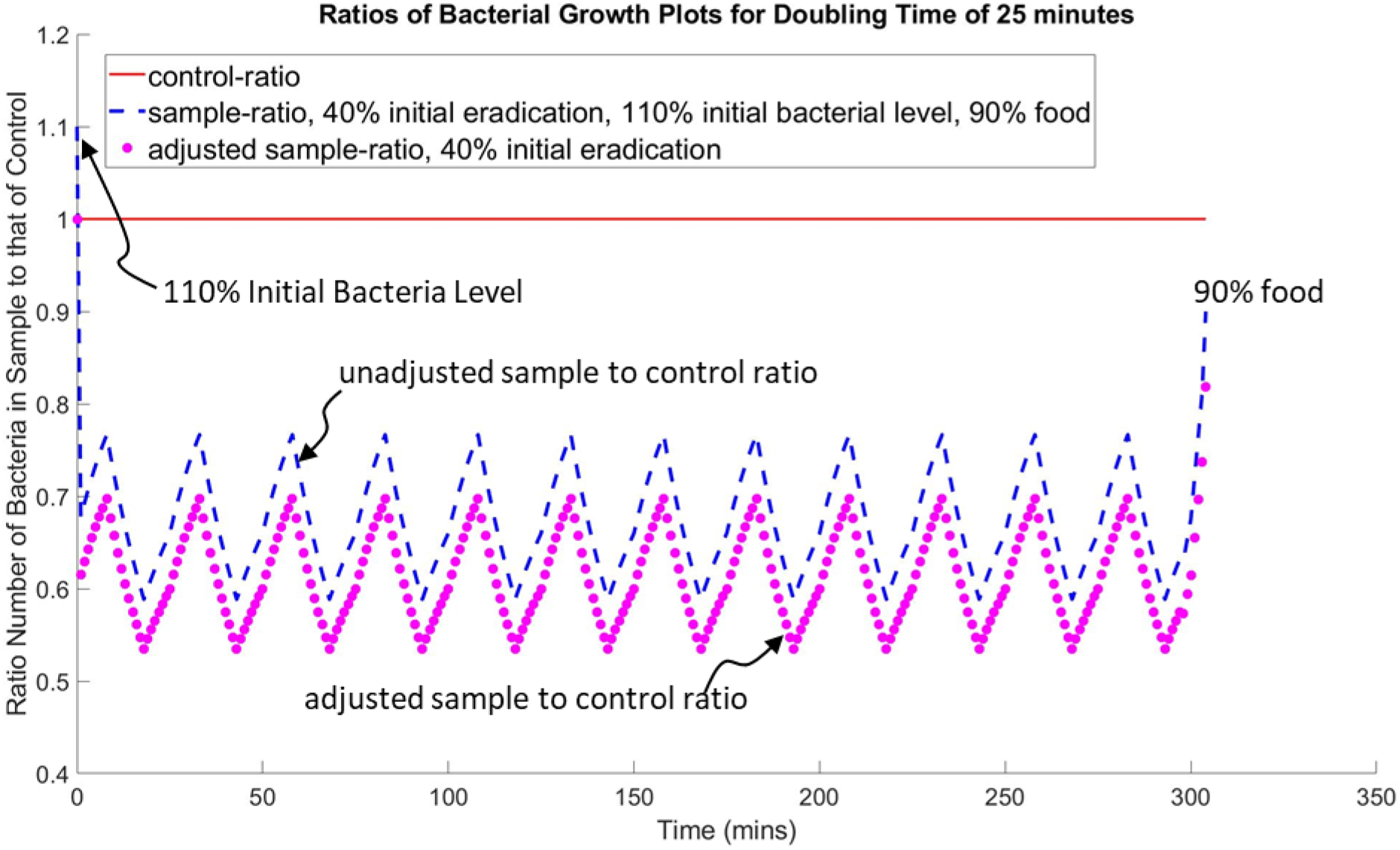
Simulated growth curve-ratio values for the various eradication levels of Figure 9

The ratio curve plots can be used to determine variations in initial bacterial levels as the ratio curves indicate different initial relative bacterial levels for different initial levels. For example, in Figure 10, the 110% initial bacterial level of the sample can be seen as the blue dashed line at the y-intercept (t=0 mins). Additionally, the ratio curves indicate different nutrient levels in the stationary phase, where 90% nutrient level, compared to the control, can be seen at time 325 minutes for the blue dashed line at 0.90. The y intercept information can be used to adjust the ratio curve such that in the exponential phase the adjusted curve oscillates about the true value of live bacterial level, or 60% for the simulated adjusted curve (purple dashed line). Adjustment is needed so that direct comparison of the sample with the control can be made from the same initial conditions when determining eradication percentage. Notice though that the 90% nutrient information is lost when artificially shifted.

### Comparison with Experiment

Data from several reported bacterial curves are extracted and evaluated in a ratio format and interpreted according to the developed model. The first of the data comes from a first paper by Deng et al. 2015, which is directed to an anti-microbial non-woven polyethylene terephthalate (PET) fabric containing nano-silver of different concentrations and discusses the effect of various concentrations on bacterial eradication. The second set of data is derived from a second paper by Sekse et al. 2012, which compares bacterial growth of E. coli to various concentrations of lactoferrin. Data from Deng 2015 and Sekse 2012, were extracted using WebPlotDigitizer-3.8^TM^. Comparison data was extracted from Figure 4A from Deng 2015, and Figure 1 of Sekse 2012.

## Results: NBF Model Analysis of Reported Experiments

Two reported experiments of bacterial eradication are examined, the first is a silver dispersion in a fabric material (Deng et al. 2015) and a second is the effectiveness of various concentrations of lactoferrin on E.coli growth (Sekse et al. 2012).Figure 11 displays growth profiles using the average of the growth values for E. coli in TSB medium, showing three general phases, lag phase, exponential phase and the stationary phase for various concentrations of silver dispersions (Deng et al. 2015). Figure 12 displays the growth curve ratios unadjusted for initial differences between bacterial levels. Initial errors in starting levels can be adjusted for by dividing the values by the initial ratio of sample to control, as indicated by eqn. 4. The adjusted ratios are displayed in Figure 13.

**Figure. 11.**
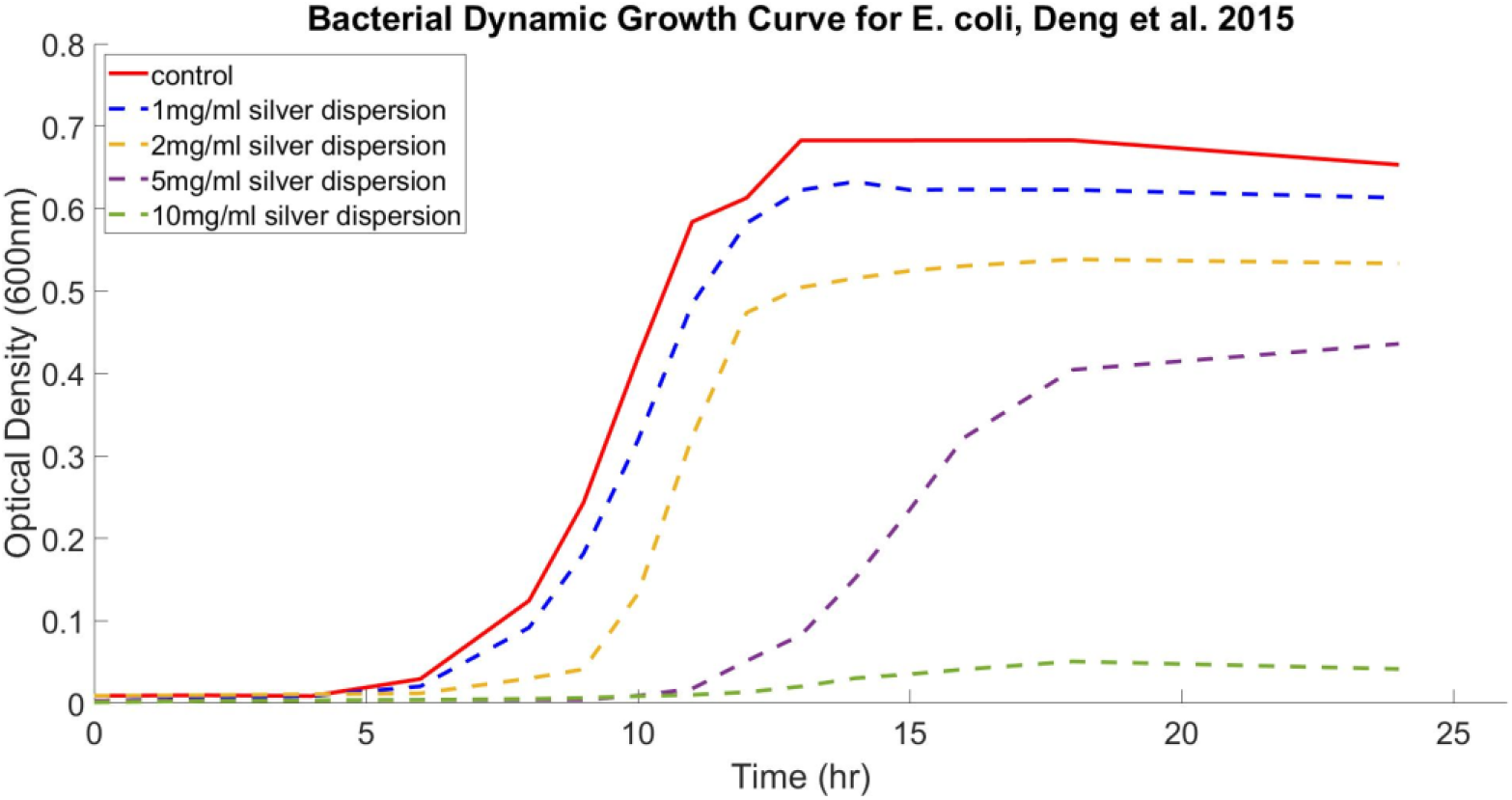
Bacterial dynamic growth curve in TSB medium with samples prepared with different concentration of silver dispersion for E. coli (Deng et al. 2015)

**Figure. 12.**
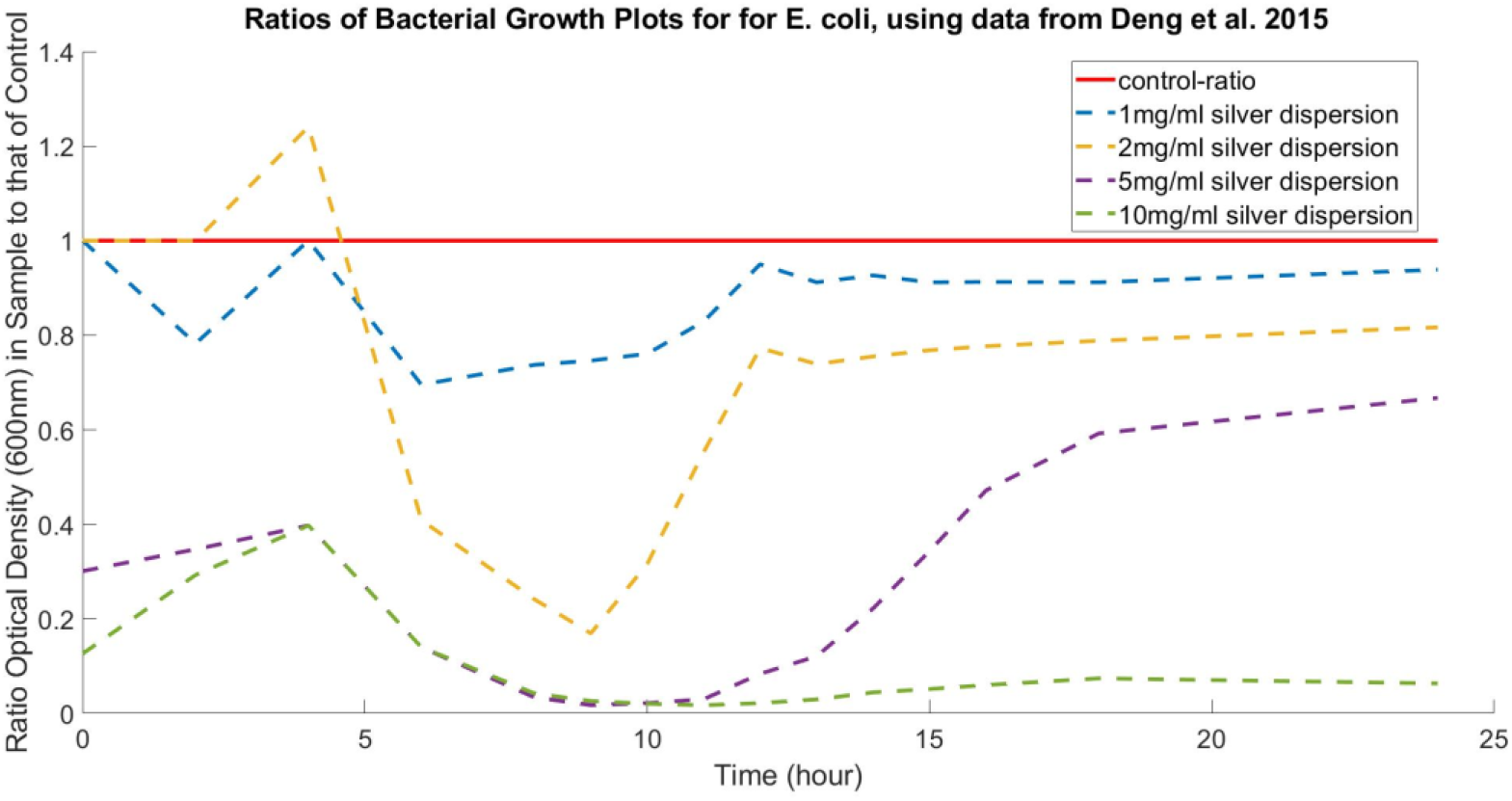
Ratios of bacterial dynamic growth curve in TSB medium with samples prepared with different concentration of silver dispersion for E. coli

**Figure. 13.**
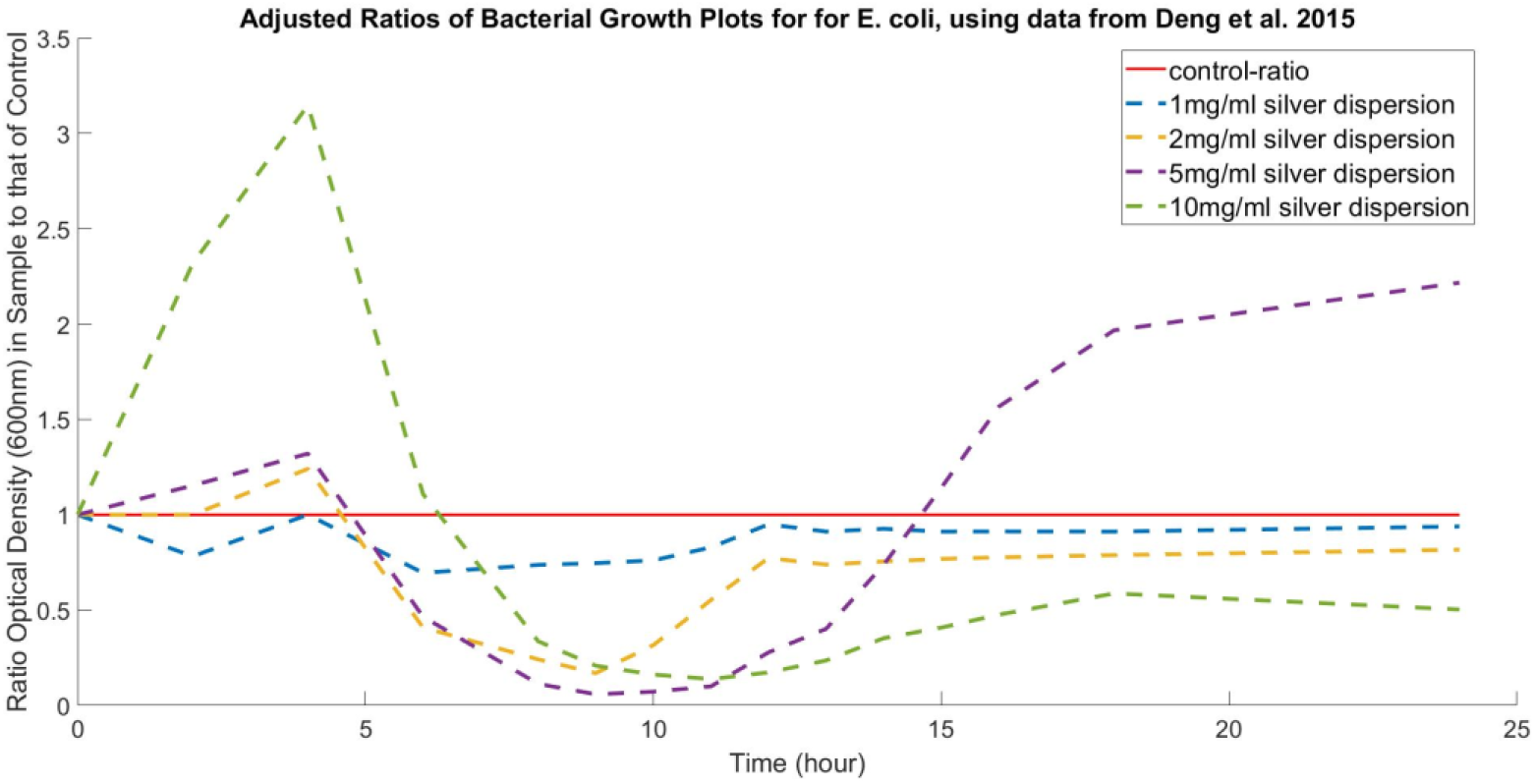
Adjusted Ratios of bacterial dynamic growth curve in TSB medium with samples prepared with different concentration of silver dispersion for E. coli

In Figure 12, an illustration of the differences in initial values, 0 hour, for the 5mg/ml and the 10mg/ml silver dispersion concentration is shown. As discussed in section 2.1, when the ratio values of sample/control of the bacterial growth, in their respective exponential stages are used, one can determine the percentage of live bacterial in the sample compared to the control. The adjusted ratio shows that eradication generally increases with increased concentration of silver dispersion. Figure 14 displays the percentage of live bacteria as a function of silver dispersion concentration derived from unadjusted and adjusted ratio curves of Figures 12 and 13 respectively. At 5mg/ml over 90% of the bacteria is eradicate with little appreciable advantage of using 10mg/ml over 5mg/ml.

**Figure. 14.**
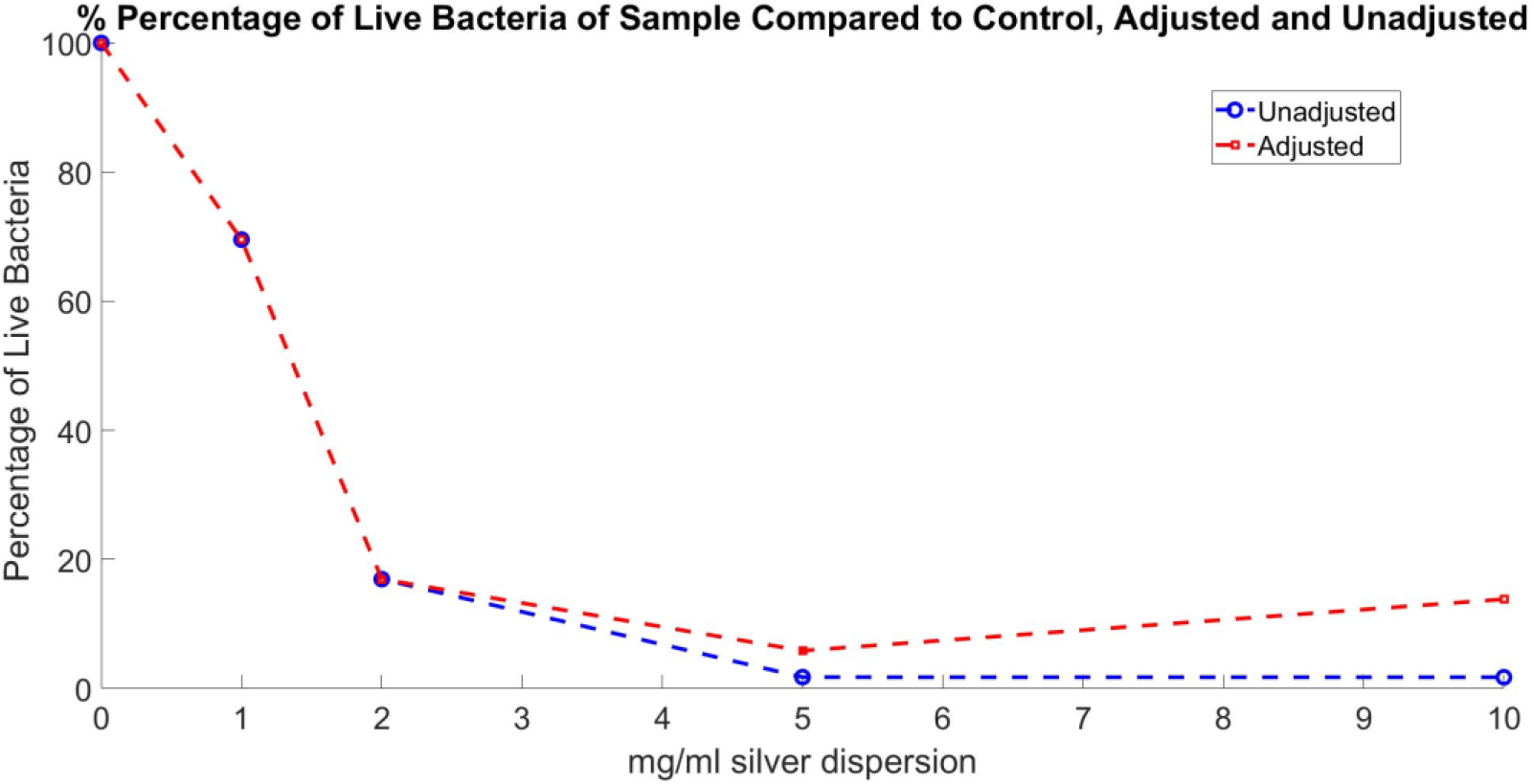
Percentage of live bacteria as a function of silver dispersion concentration for unadjusted and adjusted ration curves (Deng et al. 2015).

Figure 14 displays the derived live treated E.coli population relative to the control for various concentrations of silver dispersion. The relative live level of bacteria for the 1mg/ml, 2mg/ml, 5mg/ml, and 10mg/ml silver dispersion concentrations are 69.54%, 16.94%, 1.77%, 1.75% respectively from unadjusted ratio curves, and 69.54%, 16.94%, 5.86%, and 13.83% for adjusted ratio curves. According to the developed model, adjusted curves are needed to obtain an accurate relative percentage of live bacteria of the treated sample to control. However, since the data was extracted from plots of Deng et al. 2015, errors in point placement can result in incorrect adjustment and thus the unadjusted value and the adjusted values are presented. Figure 4A of Deng et al. 2015, from which the data for Figures 11, 12, 13, 14 were derived, does show a different start bacterial population between 5mg/ml, 10mg/ml and the other concentrations of 0mg/ml, 1mg/ml and 2mg/ml. The difference in the initial level of bacteria should be taken into account which is accomplished by using the adjusted curve of Figure 14.

Figure 15 displays growth profiles for E.coli species aEPEC, with concentrations of 0mg/ml, 3mg/ml, and 8mg/ml of antibacterial lactoferrin (Sekse et al. 2012). Figure 16 displays the ratio curve of sample to control of the curves of Figure 15 unadjusted for initial offsets, while Figure 17 displays the ratio curves for adjusted curves.

**Figure. 15.**
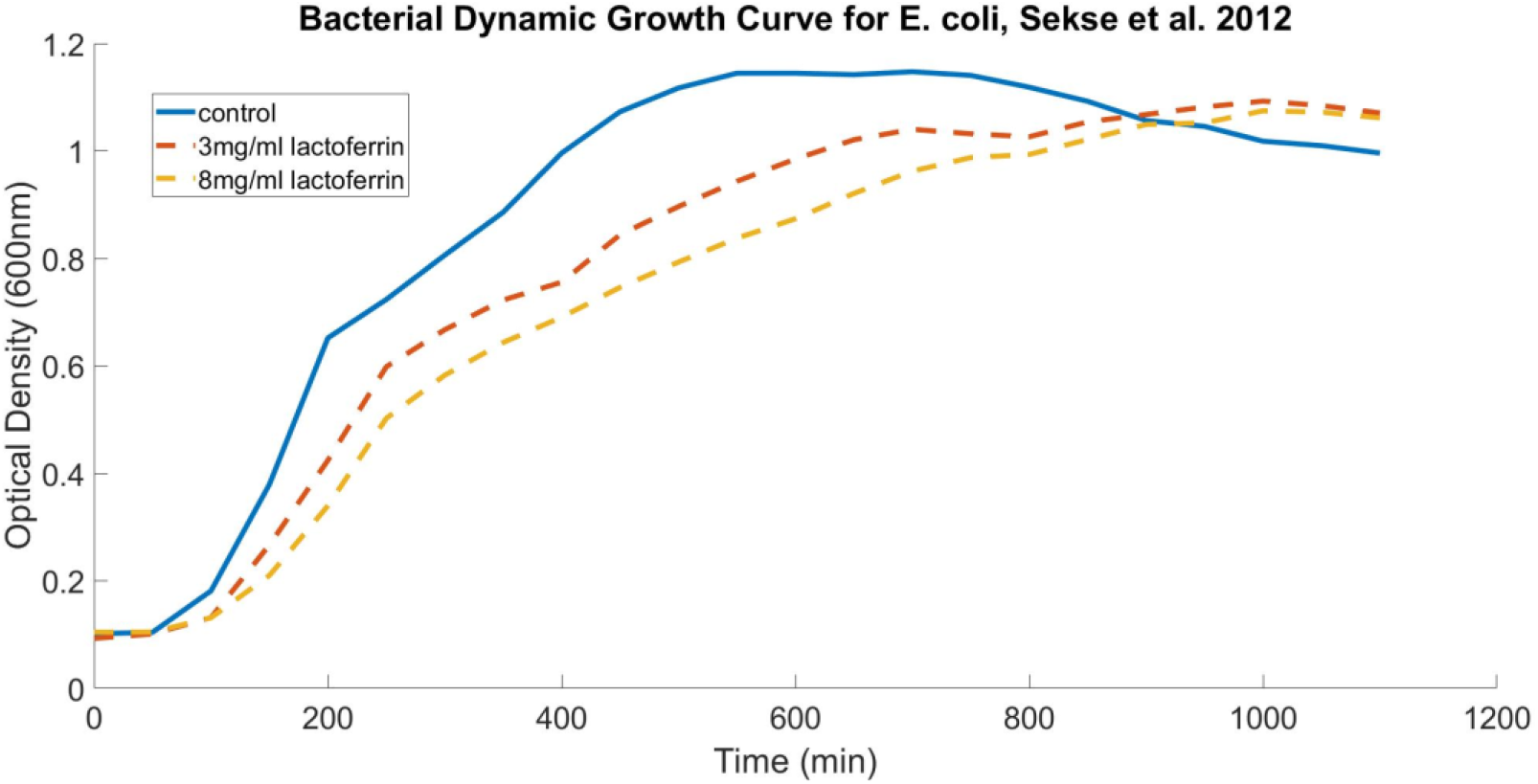
E.coli growth curves for aEPEC E.coli species exposed to 3mgl/ml and 8mg/ml lactoferrin in LB broth. (Sekse et al. 2012).

**Figure. 16.**
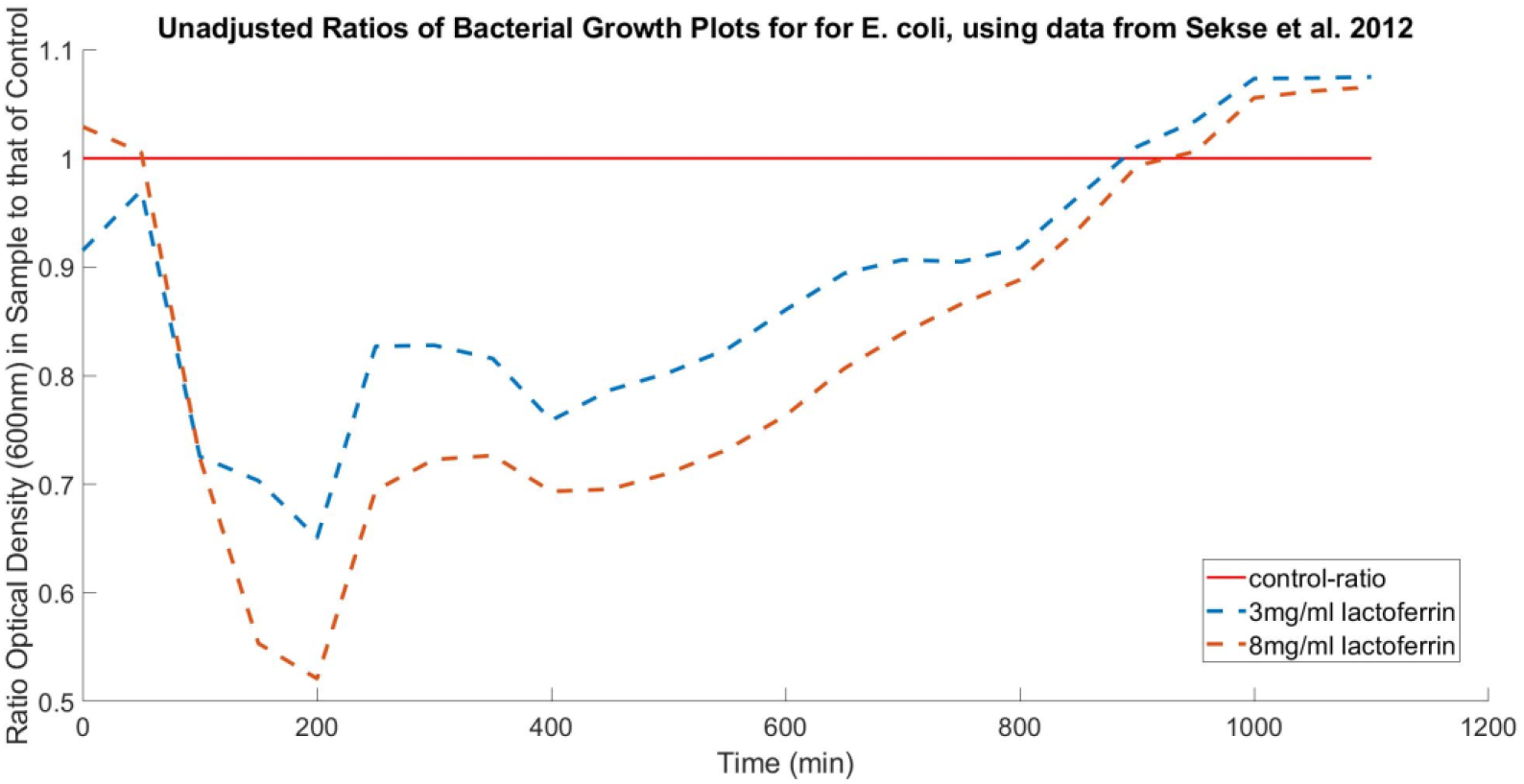
Ratios of the E.coli dynamic growth curves of Figure 15, unadjusted.

**Figure. 17.**
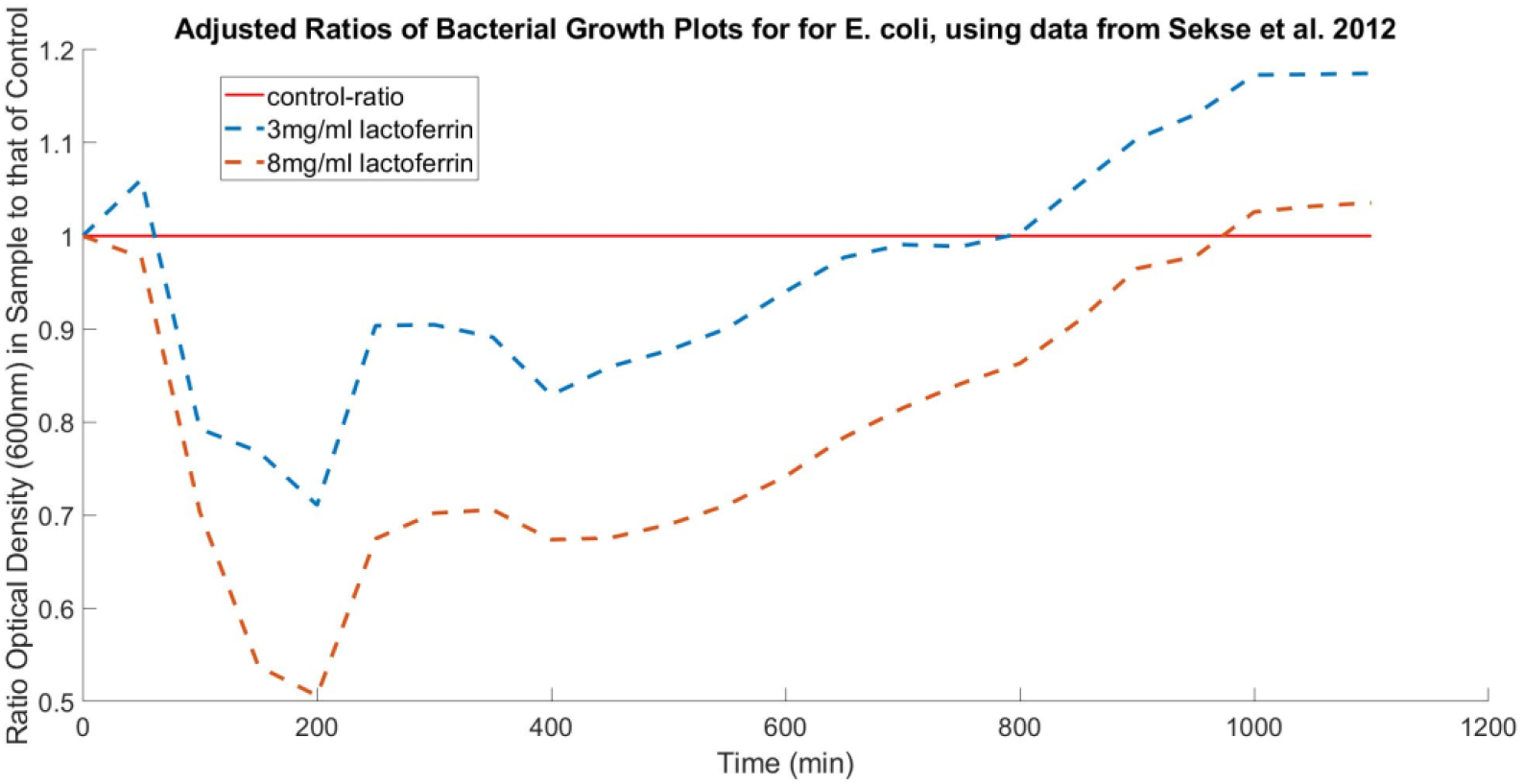
Ratios of the E.coli dynamic growth curves of Figure 15, adjusted using time = 0 hour data to adjust initial levels.

The data indicates (Figures 17 and 18) that a 3mg/ml lactoferrin has an eradication percentage of over 25% while a concentration of 8mg/ml lactoferrin has an eradication percentage of nearly 50%. Figure 18 displays the live bacterial amounts based upon the minimum of the curves of Figures 16 and 17, for unadjusted and adjusted curves. Figure 18 displays the derived live treated E.coli population relative to the control for various concentrations of lactoferrin. The relative live level of bacteria for the 3mg/ml and 8mg/ml lactoferrin concentrations are 65.08% and 52.10% respectively from unadjusted ratio curves, and 71.11% and 50.63% for adjusted ratio curves. Figure 1 of Sekse et al. 2012, from which the data for Figures 15, 16, 17, 18 were derived, shows some error in measured values where the approximate average values were point selected for data extraction at each 50 second interval.

**Figure. 18.**
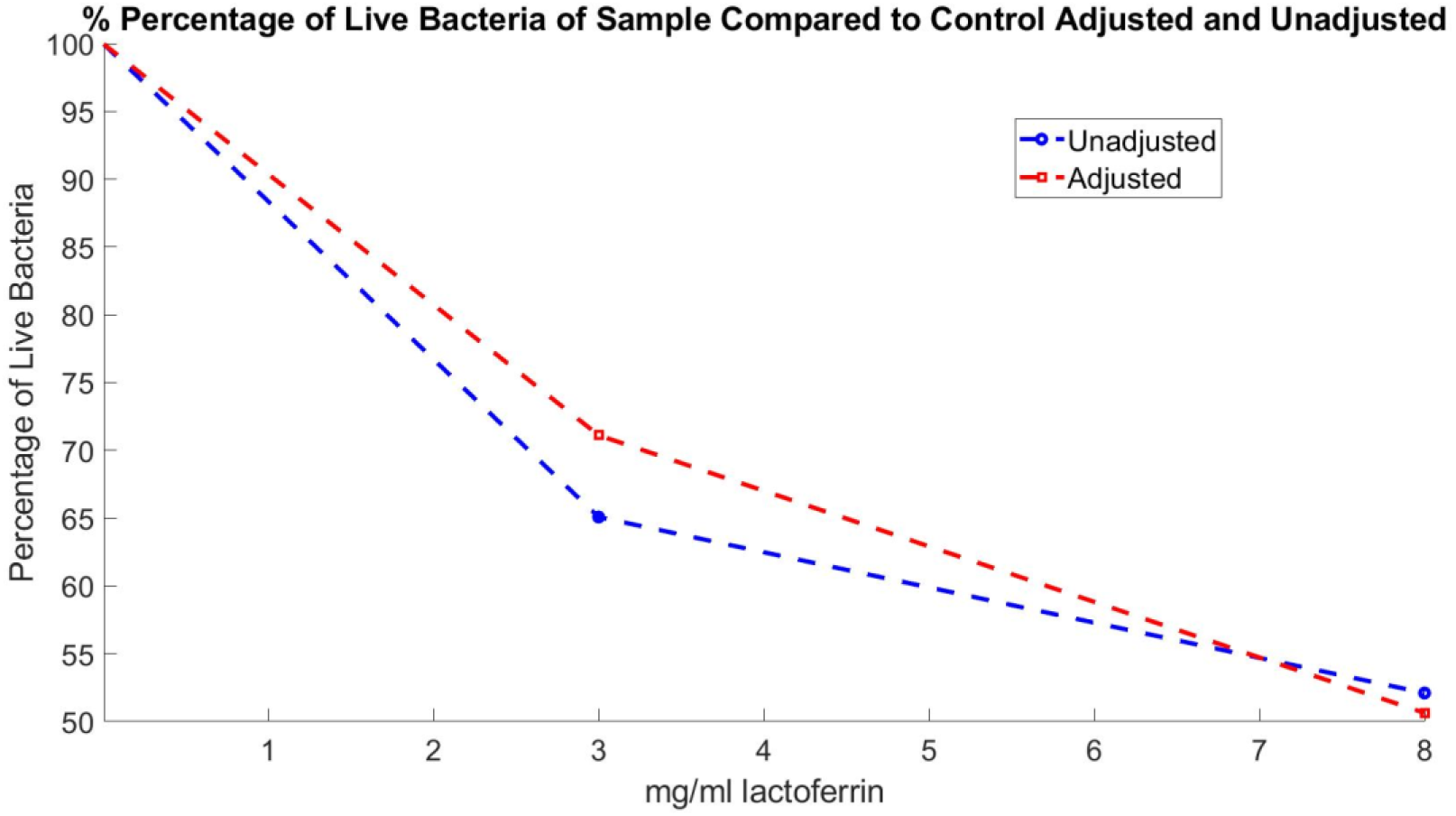
Percentage of live bacteria as a function of lactoferrin concentration for unadjusted and adjusted ration curves (Sekse et al. 2012)

## Discussion

The model above has provided a method for extracting bacterial eradication percentages using growth curve ratios (treated sample/control), a method of adjusting growth curves for variations in initial bacterial levels and method of identifying different nutrient levels between treated samples and control samples. The unadjusted ratio (treated/control) curve values at the initial time can be used to adjust treated growth curve values, which then can be used in adjusted ratio curve values to accurately obtain eradication levels in the exponential stage of growth. Accurate eradication level aides researchers in determining appropriate and exact efficacies of various treatments and dosages when comparing treated vs un-treated sample sets. For example, Figure 14 illustrates that there is not an eradication improvement when the dosage changes from 5mg/ml of silver dispersion to 10 mg/ml of silver dispersion. Additionally, the difference in beginning bacterial levels can skew the effectiveness of determining the dosage, for example in comparison between Figures 16 and 17, an unadjusted graph shows an improvement in effectiveness of less than 20% between a dosage of 3mg/ml lactoferrin and 8mg/ml of lactoferrin, while an adjusted graph illustrates an actual improvement of greater than 20%.

The methods developed using the NBF model (e.g., using the ratio of sample/control to extract information) are some of the only methods relying on indirect OD measurements, to accurately determine the effectiveness of treatments. The limitation of the methods developed using the NBF model is that they assume accurate OD measurements of both the sample and control, and close temporal measurements so that a meaningful ratios in time can be compared. The advantage of the methods developed using the NBF model are their simplicity and accuracy.

## Acknowledgments

Authors would like to acknowledge Dr. Kylene Kehn-Hall and Dr. Fernando Camelli for their support.

